# Mesendodermal cells fail to contribute to heart formation following blastocyst injection

**DOI:** 10.1101/2024.05.22.595392

**Authors:** Biyi Li, Chulan Kwon

## Abstract

Blastocyst complementation offers an opportunity for generating transplantable whole organs from donor sources. Pluripotent stem cells (PSCs) have traditionally served as the primary donor cells due to their ability to differentiate into any type of body cell. However, the use of PSCs raises ethical concerns, particularly regarding their uncontrollable differentiation potential to undesired cell lineages such as brain and germline cells. To address this issue, various strategies have been explored, including the use of genetically modified PSCs with restricted lineage potential or lineage-specified progenitor cells as donors. In this study, we tested whether nascent mesendodermal cells (MECs), which appear during early gastrulation, can be used as donor cells. To do this, we induced Bry-GFP^+^ MECs from mouse embryonic stem cells (ESCs) and introduced them into the blastocyst. While donor ESCs gave rise to various regions of embryos, including the heart, Bry-GFP^+^ MECs failed to contribute to the host embryos. This finding suggests that MECs, despite being specified from PSCs within a few days, lack the capacity to assimilate into the developing embryo.

## Introduction

As one of the earliest organs to form during embryo development, the heart plays a fundamental role in sustaining life. Despite extensive research aimed at understanding this complex organ, heart disease remains a leading cause of death worldwide. The human heart lacks the ability to repair itself after injury because cardiac muscle cells rarely renew. Consequently, numerous people, who survived a heart attack, are left with failing hearts that are typically irreversible. Thus, current cardiac regenerative studies focus on restoring the function of the damaged heart by stimulating/manipulating endogenous heart cells or delivering exogenous cells to the point of injury. While these approaches have potential in treating or preventing incomprehensive heart disease, heart transplantation remains the gold standard for treating patients with advanced heart failure. However, the scarcity of donor organs and the risk of immune rejection underscore the urgent need for alternative therapeutic approaches.

Blastocyst complementation has emerged as a promising approach for in vivo generation of donor-derived organs. Pluripotent stem cells (PSCs) can be grafted into blastocyst-stage embryos through blastocyst injection. This technique is often used to generate chimeric mice for different purposes (Gardner, 1968; Shenje et al., 2014). Blastocysts can tolerate foreign PSCs and provide the environment they need for survival and development. By introducing donor PSCs into genetically modified host embryos incapable of forming specific organs, blastocyst complementation enables the growth of complete organs with intricate 3D architectures. Moreover, these organs predominantly comprise donor-derived cells, significantly reducing the risk of rejection. Blastocyst complementation thus holds tremendous potential for revolutionizing organ transplantation by offering a route to generating rejection-free organs.

In addition to successful generation of various organs such as the pancreas, kidney, and lungs reported in previous studies (Kobayashi et al., 2010; Mori et al., 2019; Usui et al., 2012), recent success has been achieved in generating more complex organs, such as the heart (Coppiello et al., 2023). Coppiello et al. utilized Cre mediated toxin expression to delete cardiac and vascular lineages in mouse embryos, followed by subsequent rescue through injection of mouse or rat PSCs for intraspecific and interspecific complementation. In mouse PSC complemented embryos, they observed functional beating heart composed of cardiomyocytes (CMs) and endothelial cells (ECs) that are entirely derived from donor cells. In contrast to mouse PSC rescued embryos which lived to adulthood, rat PSC complemented embryos only survived to E10.5. These findings suggest the presence of interspecies barriers that impede efficient complementation and development beyond early embryonic stages.

Several obstacles such as donor-host competition and interspecies barrier need to be overcome to increase the efficiency of donor cell delivery (Mori et al., 2019). Moreover, studies involving PSCs are accompanied with ethical considerations, particularly in the context of generating human-animal chimeras. The injected human stem cells can differentiate into various cell types, raising concerns about their potential contribution to organ formation beyond the intended target, such as the brain and gonads (Kobayashi et al., 2010). Various approaches can be taken to ensure this process is safer and more controllable, including the use of genetically modified cell lines or lineage-specified donor cells. Hashimoto et al. demonstrated the feasibility of using genetically modified cell lines by injecting mouse PSCs with Prdm14 and Otx2 knockout into Pdx1 deficient embryos, thereby preventing donor cell contribution to brain and gametes (Hashimoto et al., 2019). Additionally, while previous studies have highlighted the negative impact of developmental stage differences between donor and host on successful complementation (Huang et al., 2012), certain types of PSCs retain their pluripotent potential and exhibit the ability to engraft and contribute to organ formation (Clarke et al., 2000; Geiger et al., 1998).Therefore, our study aimed to investigate the feasibility of utilizing lineage-specific donor cells for blastocyst complementation. Specifically, we explored whether mesoendodermal cells (MECs) derived from mouse embryonic stem cells (ESCs) are competent to contribute to blastocyst chimeras. Our results suggest that lineage-specified cells may not possess the same potential as ESCs for blastocyst complementation.

## Results

### MECs are generated through mesoderm differentiation of ESCs

During embryonic development, cardiac lineages originate from the precardiac mesoderm (Galdos et al., 2017). Assuming the importance of minimizing the developmental stage differences between donor cells and the inner cell mass of blastocysts, we chose to generate precardiac mesoderm cells, which may facilitate efficient integration into the blastocyst. Given that Brachyury marks nascent MECs (Herrmann, 1995), we utilized a Brachyury(T) GFP reporter mouse ESC line (referred to as TGFP ESC below). Brachyury^+^ MECs were induced through mesoderm differentiation as done previously (Andersen et al., 2018; Cheng et al., 2013). Among the Brachyury^+^ mesendodermal population, platelet-derived growth factor receptor alpha (PdgfR-α) and fetal liver kinase 1 (Flk-1) co-express on the surface of early cardiac progenitor cells (Hirata et al., 2007). Therefore, we utilized these markers for fluorescence-activated cell sorting (FACS) isolation of target MECs that will give rise to the cardiac lineage.

In order to trace the donor cells following injection into the host blastocysts and assess their contribution to embryo development and heart formation, we tagged the TGFP ESC line with red fluorescent protein mCherry through lentiviral transduction as described in the method section. Subsequently, five mCherry-tagged TGFP clones were isolated and subjected to flow cytometry analysis to evaluate RFP expression levels. We selected a pure clone with stable RFP expression (referred to as TGFP-RFP ESC below) for downstream differentiation experiments.

Next, we optimized the mesoderm differentiation condition of TGFP-RFP ESC line following an embryoid body (EB) based differentiation protocol as described in the method section. Briefly, we supplemented the culture medium with CHIR and BMP4 to activate WNT signaling and TGF-β signaling for mesoderm induction. On day 3, we observed GFP expression throughout the whole EB (Figure 1A), indicating successful generation of Brachyury^+^ mesendoderm. GFP expression became localized to one side of the EB on day 4 and diminished at around day 6. On day 7, EBs displayed irregular shapes and initiated spontaneous beating, indicative of successful differentiation into cardiomyocytes. Notably, RFP expression was stable across the entire differentiation process (Figure 1A). The percentage of cells expressing mesendoderm marker GFP and PdgfR-α were analyzed using flow cytometry (Figure 1B) on day 3 and day 4. The scatter plot (Figure 1C) summarized the percentage change of GFP^+^/PdgfR-α^+^ cells with varying BMP4 concentration. Specifically, lower concentrations of BMP4 (0 to 0.5 ng/ml) resulted in increased percentages of MECs, whereas higher concentrations led to a decrease in this population. Moreover, we observed a higher percentage of PdgfR-α^+^ cells on day 4 compared to day 3, suggesting majority of cardiac progenitor cells were specified on day 4. However, an average percentage of 20-30% GFP^+^/PdgfR-α^+^ cells on day 3 is sufficient for cell sorting and blastocyst injection. To minimize the developmental stage difference and utilize donor cells at the earliest possible stage, we performed cell sorting on day 3. Our results demonstrated that with the addition of CHIR and BMP4, TGFP-RFP ESCs are successfully differentiated into MECs for cell sorting and injection.

**Figure 1:**
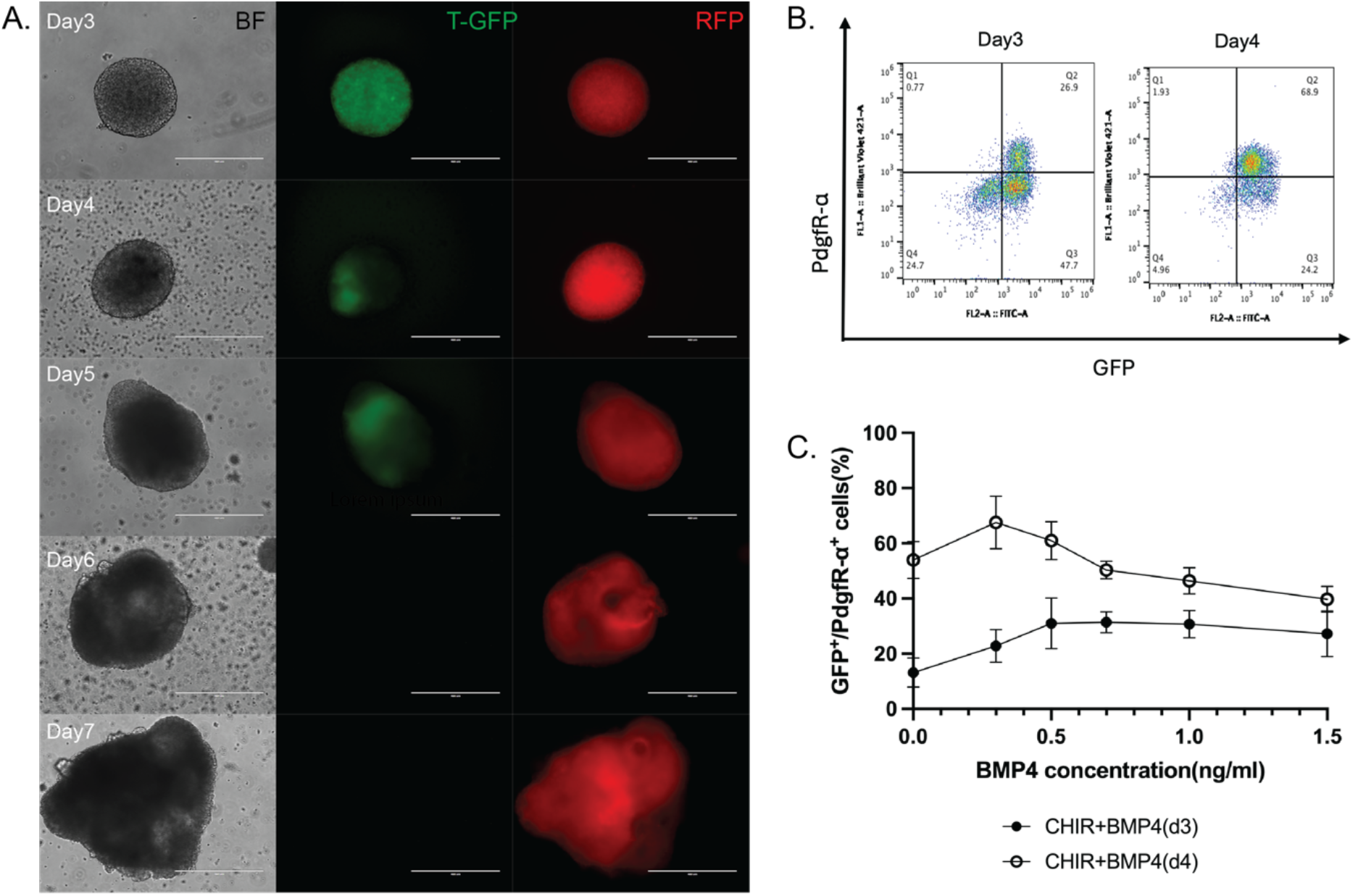
Generating MECs from TGFP-RFP mESCs. (A) Day 3 to day 7 TGFP-RFP EBs induced with CHIR& BMP4 under BF, GFP, RFP channel. T-GFP start to express around day 3 and diminish on day 6. (B) Flow cytometry analysis of day 3&4 TGFP-RFP EBs for GFP and PdgfR-α expression. The x-axis represents the GFP signal, and the y-axis represents the PdgfR-α signal. Region Q2 shows the population that is GFP^+^/PdgfR-α^+^. (C) Quantification of percentage of GFP^+^/PdgfR-α^+^ MECs on day3&4 induced with different BMP4 concentration. N=4, scale bar=400um.

### MECs injected into E3.5 blastocysts do not contribute to heart formation

After inducing differentiation of TGFP-RFP ESCs into MECs using 3μM CHIR and 0.3 ng/ml BMP4, we isolated GFP^+^/PdgfR-α^+^ population on day 3 and injected them into E3.5 wildtype mouse blastocysts. We performed 2 injections. In the first injection, 12-15 cells were injected into E3.5 wildtype mouse blastocysts and 10 injected blastocysts were transferred to 1 wildtype surrogate female mouse. In the second injection, 44 injected blastocysts were transferred to 4 wildtype surrogate female mice. Examination of E9.5 embryos revealed no RFP signals in the heart regions of the injected embryos (Figure 2B).

**Figure 2:**
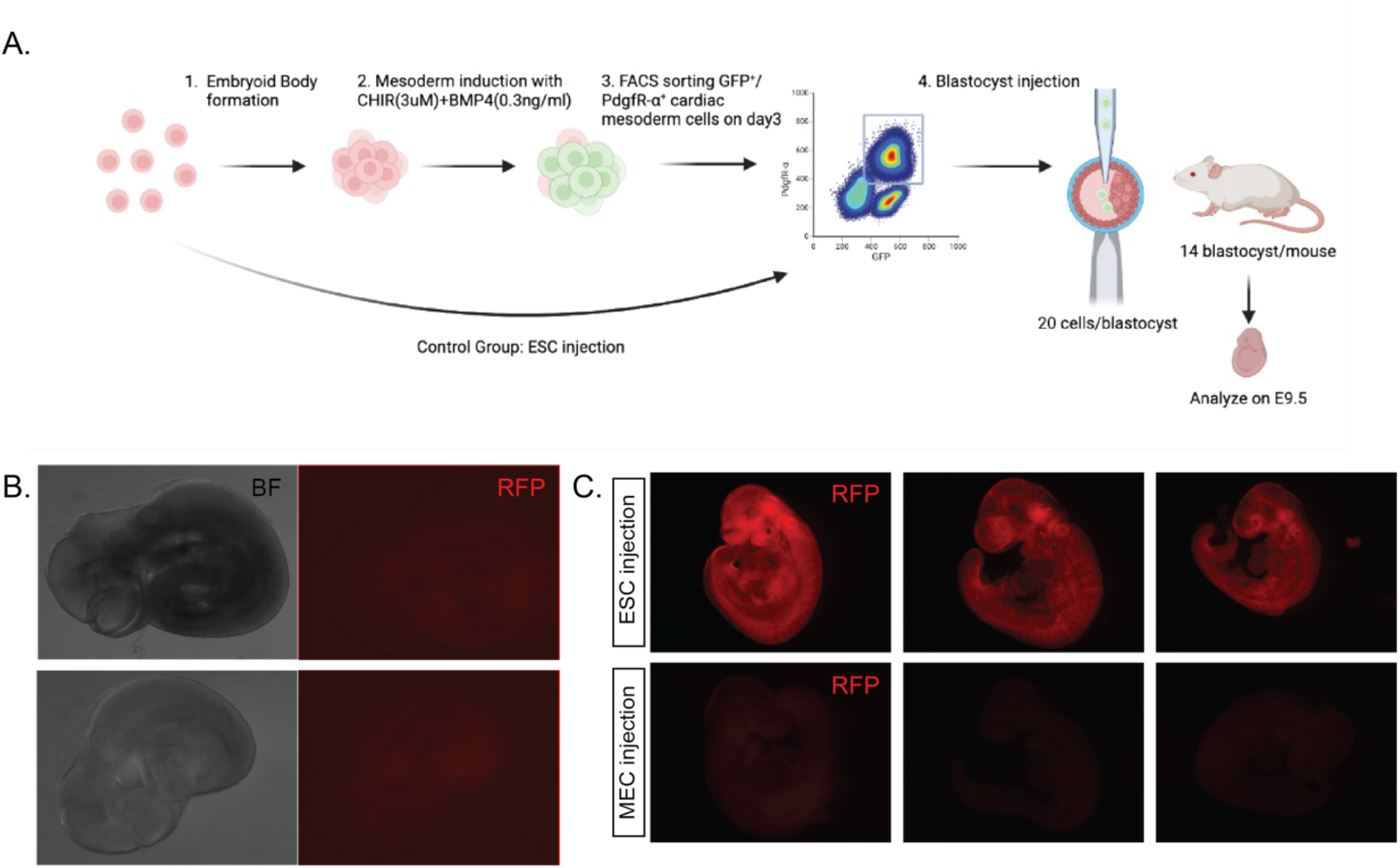
Blastocyst injection of ESCs and MECs. (A) Schematic of the blastocyst injection experimental workflow. (B) Two RFP^-^ E9.5 embryos from the first 2 injections with MECs. (C) The first row shows images of 3 RFP^+^ E9.5 embryos injected with ESCs from the third injection. The second row shows images of 3 RFP^-^ E9.5 embryos injected with MECs from the third injection.

To address the concern that the failure of donor cell engraftment was due to cell sorting induced cell stress or mis-operation during the cell preparation process, we performed a third injection and included a control group with undifferentiated mouse ESC injections. ESC injection is widely used to generate chimeric mice, and thus can be served as an indicator of whether the cell preparing process affected their ability of survival after injection. The undifferentiated mouse TGFP-RFP ESCs were subjected to the same cell preparation and sorting process as the MECs for injection. Furthermore, we increased the number of cells injected into each embryo from 12-15 cells to 20 cells to enhance engraftment potential. The experimental design is shown in Figure 2A. In this injection, 53 E3.5 blastocysts injected with MECs were transferred to 4 wildtype surrogate female mice, and 14 embryos injected with ESCs were transferred to 1 wildtype surrogate female mouse. We examined embryos at E9.5. We observed RFP signals in 7 out of 8 embryos from the ESC injected control group but none in the rest embryos injected with MECs as shown in Figure 2C. In conclusion, our preliminary findings indicate that MECs failed to integrate and contribute to embryo formation following blastocyst injection.

## Discussion

To evaluate the chimera competency of lineage-committed cells for blastocyst complementation, we conducted experiments injecting either mouse ESCs or mesendoderm-committed cells (MECs) into E3.5 mouse blastocysts and analyzed the presence of injected cells at E9.5. While we observed RFP^+^ chimeric embryos in ESC-injected group, we were unable to detect any RFP^+^ donor cells in MEC-injected embryos.

Our findings from the three injection experiments led us to conclude that MECs do not possess the same potential as ESCs in contributing to embryo development following injection into blastocysts. This disparity may be attributed to the developmental stage mismatch between the donor cells and the host embryo. Previous studies have reported that primed human PSCs (hPSCs) failed to survive after blastocyst injection (Rossant, 2008). The mesendodermal population arises during gastrulation, a process that occurs after embryo implantation (Rodaway & Patient, 2001). It is plausible that injected MECs were unable to integrate into the blastocyst-stage host embryos due to a lack of suitable niche and signaling cues for survival and differentiation. Masaki et al. demonstrated rapid death of developmentally mismatched injected cells (Masaki et al., 2016). They also showed that overexpression of anti-apoptotic gene *BCL2* in EpiSCs and Sox17^+^ endodermal cells resulted in successful blastocyst chimeras. This suggested that enhancing donor cell survival by inhibiting apoptosis can be a strategy to overcoming the developmental stage mismatch barrier.

Alternatively, modifying host embryos may enhance the likelihood of donor cell survival. Donor and host cell competition plays a crucial role in determining the donor cell integration efficiency (Zheng et al., 2021). Studies have shown that organ-deficient embryos exhibit higher complementation rates and reduced donor contribution to other organs compared to wildtype embryos. Additionally, Nishimura et al. discovered that Igf1r null mouse embryos displayed growth retardation and it gave donor cells advantages over host cells at late developmental stage which augmented the donor chimerism for interspecies blastocyst complementation (Nishimura et al., 2021). Further experiments can be carried out to investigate whether organ-deficient embryos or other types of genetically modified embryos will provide an environment with reduced competition, thereby facilitating the integration of lineage-committed donor cells.

In summary, our results demonstrate that developmental stage mismatch remains a significant barrier to the utilization of lineage-specified progenitor cells as donor cells. While blastocyst complementation holds great promise, understanding and overcoming these developmental stage barriers are crucial for its future development. Our study highlights the importance of further research focusing on donor cell selection to enhance the controllability and safety of blastocyst complementation.

## Methods

### Labeling TGFP mESCs with RFP

LV-CAG-mCherry from SignaGen was used for lentiviral transduction as done (Cho et al., 2017). TGFP mESCs were seeded into three wells of a 6-well plate with gelatin coating in 2ml of 2i media per well. Different volumes of lentivirus (1.5μl, 2μl, 3μl) were added to each well accordingly. After 24 hours of incubation, media with the virus was aspirated and replaced with 2ml of 2i media containing 4μl of puromycin per well to start antibiotic selection. Cells from the three wells were trypsinized when confluent and expanded into a 15cm cell culture dish. Single colonies with uniform RFP expression were picked and expanded in a 96-well plate. Each well was transferred to a 10cm petri dish for further expansion and colony picking. Media containing puromycin was changed regularly during the selection process.

### Mouse ESC mesoderm differentiation

TGFP-RFP ESCs were cultured on a 10% gelatin-coated T-25 flask in 5 ml of 2i+LIF media before differentiation. Media was changed to LIF only media when cells reached 60%-70% confluency. After 24 hours in LIF media, cells were trypsinized, washed with PBS, and resuspended in Serum-Free Differentiation (SFD) media. Cells were then transferred to a 10cm petri dish with a seeding density of 75k cells/ml for embryoid body (EB) formation. After 48 hours, EBs were collected and transferred to a 6-well Ultra-Low Attachment Plate in SFD media.

CHIR99021& BMP4 were used for mesodermal induction. Each well was treated with 3μM CHIR99021, and BMP4 of different concentrations (0, 0.3, 0.5, 0.7, 1, 1.5 ng/ml) for 24 hours. Media was changed every two days after removing cytokines.

### Sample preparation for flow cytometry analysis

To prepare samples for flow cytometry analysis, EBs were dissociated into single cells and stained with different antibodies.

For dissociation, EBs were collected on day3 into 96-well U bottom plates and centrifuged at 800 rpm for 3 minutes. The supernatant was discarded, and 100ul of trypsin was added to each well. The plate was then incubated at 37°C for 3 minutes. Each well was added 50ul FBS to neutralize the reaction and resuspended to make sure the EBs are dissociated into single cells. The plate was then centrifuged at 1100 rpm for 3 minutes. The supernatant was discarded, and dissociated cells were resuspended in PBS and ready for analysis.

For PdgfR-α staining, BD Horizen™ BV421 Rat Anti-Mouse CD140A antibody (Cat: 562774)(1:500) was added after the dissociation process. Cells were washed with PBS for three times before analysis.

### Cell sorting and blastocyst injection

Day 3 GFP^+^/ PdgfR-α^+^ cardiac progenitor cells were dissociated, sorted using the Sony cell sorter (SH800), and resuspended in M2 injection media. Blastocyst injections were performed by the transgenic core at Johns Hopkins University as done previously (Gibbs et al., 2018; Shenje et al., 2014).

### Embryo dissections

E9.5 embryos were collected, cleaned in PBS containing calcium and magnesium, and imaged under the dissection microscope for RFP signal.

## Acknowledgements

The authors thank Kwon laboratory members for critical reading and discussion. This study was supported by awards from PRMRP/DoD (W81XWH-20-1-0078), NHLBI/NIH (R01HL156947), and AHA (969345).

## Disclosers

None.

